# Alterations in the Ultrasonic Vocalization Sequences in pups of an autism spectrum disorder mouse model: A longitudinal study over age and sex

**DOI:** 10.1101/2023.12.24.573249

**Authors:** Swapna Agarwalla, Yuvarani Selvarajan, Sharba Bandyopadhyay

## Abstract

Social communication deficit is a hallmark of autism spectrum disorders (ASDs). Mouse ultrasonic-vocalizations (USVs), with communicative significance, are extensively used to probe vocalization-based social communication impairment. Despite the predictable nature of mouse USVs, very few studies have taken advantage of the same. The current work explores USV pup-isolation-call (PIC) features and alterations in structural content of predictive PIC sequences of the well-established *in-utero* valproic-acid (VPA) exposure-based ASDs model. Our study shows that along with call features, even higher-order USV structures undergo alterations in the ASDs model at all developmental ages and sexes. Confirming prior observations, we found reduced call rates and durations, as well as heightened peak frequencies in ASD model pups. Our data also highlights trends in call features, syllable composition, and transitions across sexes and age. The ASD female mice exhibited higher within group heterogeneity in syllable composition and transition over age compared to ASD males or typically developing males and females. Analysis of sequences of USVs emitted by pups using mutual information between syllables at different positions revealed that dependencies between syllables were higher in typically developing mice of both sexes compared to ASD model pups. In brief, we found that PICs call features were altered in VPA mouse models both for male and female pups and their vocalizations lack the complex syllable sequence order emitted by typically developing ones. Our studies will help establish and further investigate ASD mouse models to get a clearer picture of abnormalities related to social communication deficits over sexes and age.

## Introduction

Autism Spectrum Disorder (ASDs) is a neurodevelopmental disorder characterized by various deficits, one of the common and earliest symptoms of which is the impairment in speech development (Mody and Belliveau, 2013). Mouse Ultrasonic Vocalization Sequences (USVs) and human speech both involve vocalizations that are of communicative significance, but they have distinct characteristics and complexity levels, making them difficult to compare (Portfors and Perkel, 2014). Nevertheless, at the rudimentary level, any alterations in the structure of the USVs emitted could serve as readout of communicative deficits in laboratory conditions. Thus, pup USVs emitted have been utilized to identify ASDs-like behaviors, which are not easily measurable with other behavioral indices during early developmental age (Semple et al. 2013; Silverman et al. 2010).

Among the various ASD models, *in-utero* exposure to Valproic acid (VPA) is well-established (Moore et al. 2000; Williams et al. 2001; Rasalam et al. 2005; Rodier et al. 1996). A resemblance in behavioral deficits has been observed in prenatally VPA-exposed rodents as in human offsprings (Chomiak et al. 2013). Since then, various studies have explored the pathways linked with ASDs-like behavioral impairments using the VPA model, which has a similarity in the underlying cause and the spectrum of human ASD phenotypes (Chomiak et al. 2013; Roulette et al. 2019). Some studies previously characterized autistic-like phenotypes in VPA models with either a bias for male offspring (Cheha et al. 2015; Kuo et al. 2017; Moldrich et al. 2013) or not focusing on sex specificity (Gandal et al. 2010, Mehta et al. 2011) while others have explored the same in a sex-specific way (Tartaglione et al. 2019,Tsuji et al. 2020). For instance, several studies have observed altered call features in VPA mice at different developmental stages, with varying effects across sexes. Decreases in call rate have been observed in combined mouse samples across multiple postnatal days (p2, p5, p8, and p12; Gandal et al. 2010), in male only at p8 (Kuo et al. 2017; Moldrich et al. 2013) and p12 (Cheha et al. 2015). Along with the effect on call rate in male at p8, reduced duration and peak frequency were also observed (Kuo et al. 2017). Studies have also been done in a sex-specific way by considering mice from p4-p12 and the alteration was observed only in females at p10 that emitted shorter calls, but no significant effects were observed on other postnatal days (Tartaglione et al. 2019). Another study (Tsuji et al. 2020) investigated VPA mice of both sexes from p3 to p14 and found lower call rates in both sexes only at p11 but no effect on duration. Again, alteration in syllable composition has been reported only at p8 (Cheha et al. 2015, Moldrich et al. 2013), p12 (Cheha et al. 2015 and p11 (Tsuji et al. 2020), but a longitudinal study over developmental age in VPA mice is missing. While a limited number of studies have assessed the predictability of mouse USV sequences in control pups (Grimsley et al. 2011) and adults (Agarwalla et al. 2023; Grimsley et al. 2011). Only one study to the best of our knowledge has reported altered USV sequences for a genetic mouse model of ASDs at p5 (Agarwalla et al. 2020). By focusing on call features along with vocalization structures that are affected over sex and age, we can more accurately characterize autism during the neonatal phase.

Our work focuses on the differences between typically developing (TD) and VPA pups over developmental age, considering the USV acoustic features, the composition of syllables, and also leveraging the predictiveness and presence of higher-order vocal structures well known in mice by tracking high-order informative sequences (Agarwalla et al. 2020; Agarwalla et al. 2023). Our findings corroborate prior studies, indicating a general decrease in call rate (Cheha et al. 2015; Kuo et al. 2017; Moldrich et al. 2013; Gandal et al. 2010; Tsuji et al. 2020), heightened peak frequency (Kuo et al. 2017), and shorter call duration (Kuo et al. 2017) in VPA pups. Alterations in higher-order vocal structures and dependencies between syllables were notably pronounced in VPA pups regardless of sex and age. We observed alterations and lack of the complex syllable sequence order in USV sequences in VPA pups, suggesting the potential for these sequences to also serve as an indicator of early socio-communicative deficits in ASD models. Thus, our study opens avenues to utilize structural analyses of neonatal vocalizations along with call features to assess early socio-communicative deficits in ASD models across sex and age.

## Methods and Materials

All methods were carried out in accordance with relevant guidelines and regulations. The study design is explained by the schematic shown in Fig. 1A. The parameters assessed included call rate, mean peak frequency, call duration, frequency of occurrence of each syllable type, transition probability of syllables, Mutual Information (MI) content between the syllables in the starting and the other successive positions and tracking of informative sequences. The statistical analysis included RM ANOVA followed by post hoc Tukey multi comparison to compare TD and VPA groups over age for male and female pups. The significance of probability distributions, distance between them computed using Kullback Leibler Divergence (KLD) and MI content was estimated by bootstrapping and determining the 95% CI compared to the scrambled data. The vocalization recordings were performed from pups with age group p5, p7, p9, p11 and p13 and the sex was identified. For all the results, the mean+/-sem or the distributions has been shown in the plots.

**Figure 1.**
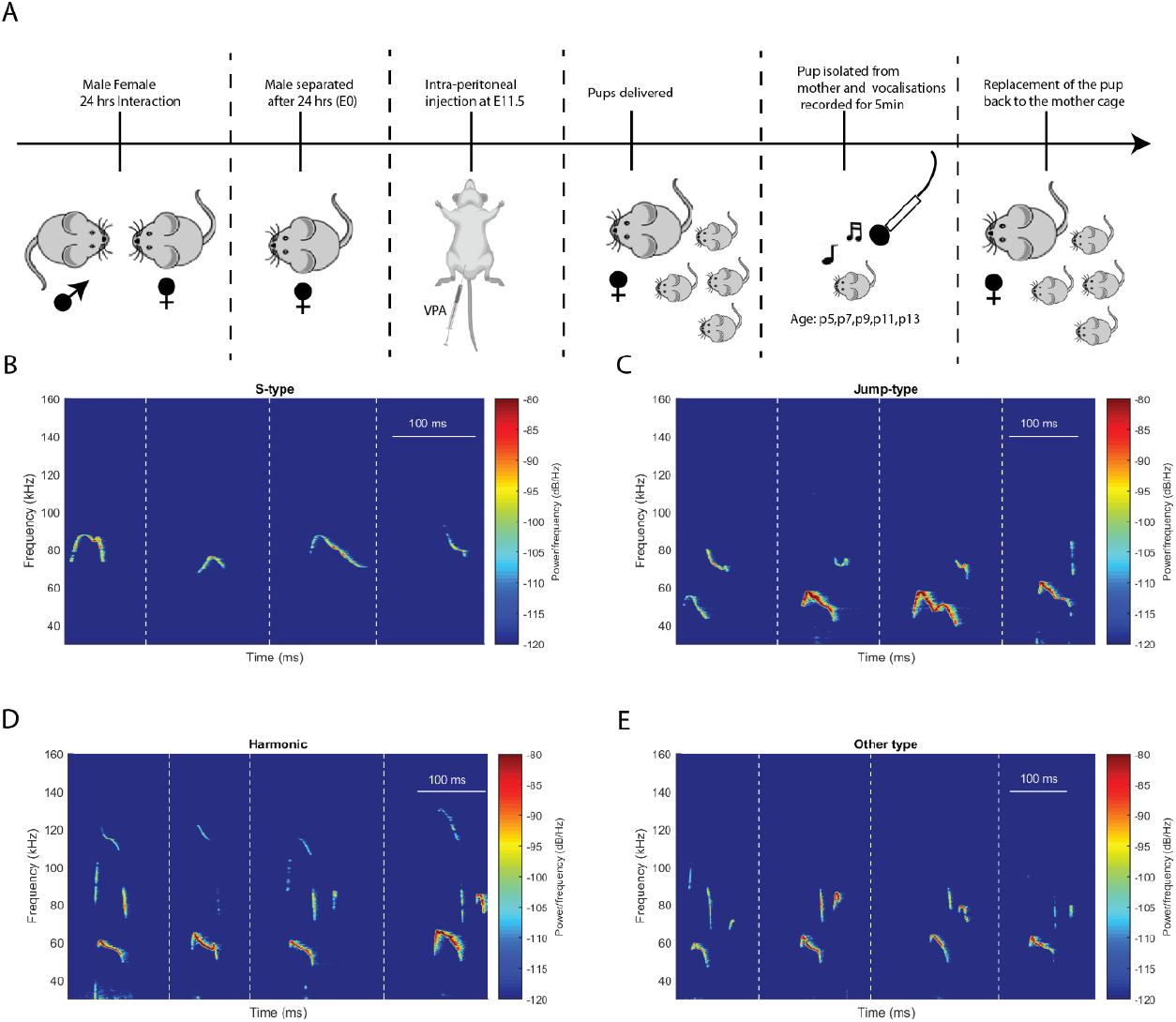
A) Schematic of the generation of the VPA model and recording of pup-isolation-calls (PICs). Representative spectrogram of syllables types B) S-type C) Jump-type D) Harmonic E) Other types. Four examples are shown for each of the syllable types (A-E). Dashed vertical white lines show different examples of a syllable category. Scale bar shown on the top right of each figure.

### Animals

All procedures used in the study pertaining to animals were approved by the Institutional Animal Ethics Committee (IAEC) of the Indian Institute of Technology Kharagpur. C57BL6/J strain mice, *mus musculus* (age and sex identified in individual cases) used in the study were reared under a 12/12 h light/dark cycle at a temperature of 22–25°C with *ad libitum* access to food and water. Vocalization recordings were performed on a total of 15 pups, comprising 8 males (Mp) and 7 females (Fp) pups. In order to remove potential litter bias, 15 pups were randomly chosen from 3 different litters from 3 distinct breeding pairs. Each litter contributed at least 4 pups (2 male and 2 female) to the recordings, encompassing the age range between p5 and p13 (inclusive of odd-numbered day age groups within this range). ASD mouse models were created by administering valproic acid, an anti-epileptic drug (details given in the VPA treatment section of materials and methods). Subsequently, recordings were conducted on 12 pups, consisting of 6 males and 6 females from three distinct sets of parents, with pups chosen randomly from each litter. None of the animals and vocalization waveforms were excluded from the analysis. For VPA treatment females were randomly chosen not following any feature or exhibiting a particular behavior. Blinding was not needed as the process of analysis of syllables was automated except for setting the threshold above the noise floor which was confirmed by the investigators. The saline group comprised a total of 8 pups selected from three distinct sets of parents, with pups chosen randomly from each litter.

### VPA Treatment

An adult female was checked for the estrus cycle since the chances of conception are pretty high during the estrus phase. The estrus state of the female was determined by visually inspecting her external genitalia. Generally, if the vaginal opening appeared swollen, pink, and moist, the female was considered to be in the estrus cycle. The male and the female (in estrus) were allowed to interact for a time span of 24h (time and date noted), following which the male was separated, and the female was checked for a vaginal plug, the day considered to be the gestational day ‘0’ or embryonic day ‘0’ (E0). For creating a VPA-induced ASDs mouse model, the timed pregnant female received a single intraperitoneal injection of a freshly prepared sodium salt of VPA (NaVPA-Sigma_Life Science, P4543) at a concentration of 400 mg/kg of mouse body weight dissolved in 1X Phosphate Buffered Saline (Gibco_70011-044) with a dosing volume of 100uL on the gestational day E11.5-E12.5^20^.The Saline group had the same volume of saline injected intraperitoneally instead of VPA during the same time. The TD group had undergone no such injection procedure. The mice were housed individually (left undisturbed until the pups got delivered) and allowed to raise their litter. All the VPA-based experiments were carried out on these delivered pups. The pup isolation calls (PICs) from the delivered pups were recorded from p5-p15 (on odd alternative days).

### Syllables: Identification and classification

Recorded sound signals in each .wav file were bandpass filtered, removing frequencies below 30 kHz and above 160 kHz for pups. For each chunk, a short term Fourier transform was calculated using a Hanning window of length 1024 and an overlap of 75%. Syllable occurrences were identified based on power concentration and syllable classification was based on the presence or absence of a distinctive feature, pitch jump^21^. The methods followed for syllable identification were similar to those reported in Agarwalla et al. 2020 and were done in MATLAB. In brief, for syllable identification, peaks were detected using a peak detection method from the power spectrum of the vocalization recording that compares each data element to its neighboring values. If an element was larger than both its neighbors, it was identified as a local peak. These local peaks were tracked until their values fall below a threshold (mean + 0.01 * STD). This process continued iteratively until all peaks were identified and stored as syllables. Call features like call rate, the average number of calls emitted by a pup per minute; the duration of the syllable and peak frequency of the syllable i.e; the lowest frequency with the highest power concentration were quantified. Each vocalization contributed to the mean for each animal; each individual’s mean was then averaged to obtain the group mean for the call features. Four categories of syllable types were considered, namely S-type (S, no discontinuity in the frequency contour, no pitch jump), Jump type (J, single jump in frequency), Harmonics (H, presence of fundamental and harmonic components), and Other type (O, more than one jump in the frequency) (Agarwalla et al. 2020; Agarwalla et al. 2023). An intersyllable interval of greater than 350ms was considered to be the beginning of a new bout (Agarwalla et al. 2020).

### Calculation of Joint Distributions and *MI* Based Dependence

For a syllable-to-syllable transition, joint distributions were estimated and cases with zero values were handled by applying Krischevsky and Trofimov (KT) correction (Krichevsky and Trofimov, 1981). Mutual information (Cover 1999) was computed based on joint distribution *P*(*X, Y*) and its marginal distributions *P*(*X*) and *P*(*Y*) as:

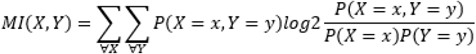

MI was calculated between the syllable in the first position with the syllable occuring in the nth position (n=1, 2, 3, …) to quantify dependency. Bootstrap (Efron and Tibshirani, 1994) was used to remove bias in the estimates and a comparison was made with the scrambled version of the original one (Agarwalla et al. 2020; Agarwalla et al. 2023). The details of the steps followed were adopted from Agarwalla et al. 2020, 2023.

### Kullback Leibler Divergence (*KLD*) between distributions

To compute *KLD* (Cover 1999) between the distributions *p* and *q* (in our case syllables produced by say two different ages (or groups) of pups, for example typically developing male at p5 and p7 taking values of different syllable types with probabilities *p*(*x*) and *q*(*x*), *x* being a syllable type, or the syllable-to-syllable transitions produced at p5 and p7 as given below

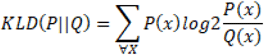

Debiasing of KLD was performed using bootstrap resampling, with significance determined by 95% confidence intervals. The details of the steps followed were adopted from Agarwalla et al. 2020, 2023. Significance is determined by was estimated by bootstrapping 1000 times and determining the 95% CI compared to the scrambled data.

### Pup Vocalisation Recording protocol

Male and female pups were individually placed on clean bedding material in a glass container (10 cm × 8 cm × 7 cm; open surface). The recording was made on postnatal days (p) p5, p7, p9, p11, and p13 for 5 minutes. After each recording, the glass container was wiped with 70% alcohol, and the bedding was changed. Acoustic signals were recorded with a free-field microphone and amplifier (1/4” microphone, Model 4939, Bruel and Kjaer, Naerum, Denmark), placed 5 cm above the pup. The acoustic signals were digitized at 375 kHz with 16-bit resolution collected with National Instruments DAQ. Recorded signal time waveforms and spectrograms were displayed in real-time on a computer with open access bioacoustics software, Ishmael.

### Tracking of Informative Sequences

The calculation involved determining the significance of syllable occurrences in various positions, based on the preceding syllables as reported by previous studies (Agarwalla et al. 2020; Agarwalla et al. 2023). For the initial syllable within a sequence, we calculated the likelihood of each syllable type occurring at the sequence’s outset. This likelihood was then compared to the probability of each syllable type occurring with equal likelihood. Syllables exhibiting a higher probability of occurrence (with 95% confidence) than the overall average were deemed significant. This process was iterated for subsequent positions, with the preceding syllable types held constant until no more statistically significant syllables were identified. Through this approach, we identified sequences that surpassed random chance, thereby revealing underlying structural patterns.

### Removal of Bias in Mutual Information Estimates and Significance Analysis

Bootstrap debiasing (Efron and Tibshirani, 1994) was utilized to mitigate bias in mutual information estimates as reported by previous studies (Agarwalla et al. 2020; Agarwalla et al. 2023).. This involved employing a resampling technique where the bootstrap dataset matched the original dataset’s element count. The aim was to rectify the bias and derive confidence intervals for the debiased estimates. To gauge significance, a comparison was made between the lower and upper confidence intervals of the ‘0’ MI estimate. This estimate was derived by deliberately scrambling the sequences, disrupting the natural transitions between syllables. This led to the estimation of ‘0’ MI and its corresponding confidence interval from the same dataset, maintaining the same syllable count and other statistics, save for the sequence syllable order.The criterion for significance was established: if the confidence intervals of MI from the original data and the scrambled data did not overlap, the MI estimate was considered statistically meaningful. This approach effectively minimized the potential (<5%) for spurious MI due to the limitations of data size and inherent variability.

## Data Availability Statement

The datasets used and/or analyzed during the current study are available from the corresponding author on reasonable request.

Computational Neuroscience Model Code Accessibility Comments for Author: Not applicable

## Results

To elucidate the differences in VPA mouse models compared to typically developing ones for both male and female pups, we initiated our investigation by examining the basic acoustic features of the syllables, namely call rate, syllable duration, and mean peak frequency. In Fig. 1A, the steps involved in generating the VPA model until the recording of pup vocalizations are depicted. The pups were isolated from their mother, and their vocalizations were recorded for 5 min (see Methods and Materials for details). In Fig. 1B-E, four example spectrograms of each syllable type are shown. Call rate, syllable duration and mean peak frequency were quantified for acoustic features. Repeated measure (RM) ANOVA was performed to assess the effect of treatment (TD or VPA) x Sex x Age. Subsequently, post hoc analysis was performed with Tukey’s multiple comparison test. The sample size used in the study containing the number of animals, litters, syllables and bouts, is tabulated in Table 1.

**Table 1:** Sample size used for the current study. The table includes numbers of animals(n), litters from different parents, syllables and bouts for different age groups in TD and VPA for male and female offsprings. Notably each litter had a minimum of 2 pups.

| **TDMp(n=8,**<br>**3 litters)** | **#syllables** | **#bouts** | **TDFp(n=7,**<br>**3 litters)** | **#syllables** | **#bouts** |
| --- | --- | --- | --- | --- | --- |
| p5 | 2443 | 494 | p5 | 2146 | 418 |
| p7 | 3252 | 713 | p7 | 3757 | 754 |
| p9 | 1929 | 499 | p9 | 2251 | 643 |
| p11 | 1270 | 379 | p11 | 2197 | 606 |
| p13 | 1254 | 401 | p13 | 884 | 386 |
| **VPAMp(n=6,**<br>**3 litters)** | **#syllables** | **#bouts** | **VPAFp(n=6,**<br>**3 litters)** | **#syllables** | **#bouts** |
| p5 | 1957 | 443 | p5 | 1845 | 400 |
| p7 | 1319 | 363 | p7 | 1237 | 373 |
| p9 | 979 | 506 | p9 | 1894 | 679 |
| p11 | 1253 | 412 | p11 | 1787 | 346 |
| p13 | 435 | 161 | p13 | 772 | 307 |

### Call rate

The average number of calls emitted by pups per minute is shown in Fig. 2. Independent of age and sex, the average call rate was lower for VPA compared to TD (RM ANOVA). A pairwise Mann-Whitney U-test revealed that VPA male pups (VPAMp) exhibited significantly fewer calls than typically developing male pups (TDMp) at p7(p<0.05) and p9(p<0.05) as shown in Fig. 2A, and for VPA female pups (VPAFp) at p7(p<0.05) indicated in Fig. 2B. A RM ANOVA reveals a significant effect of age on call rate (F (4,92) =7.3, p<0.001). However, there is no significant interaction between group types i.e;(VPA or TD) x Sex with age (F(12,92)=1.3,p>0.05). No sex specific differences were observed between the TDMp and TDFp.

**Figure 2.**
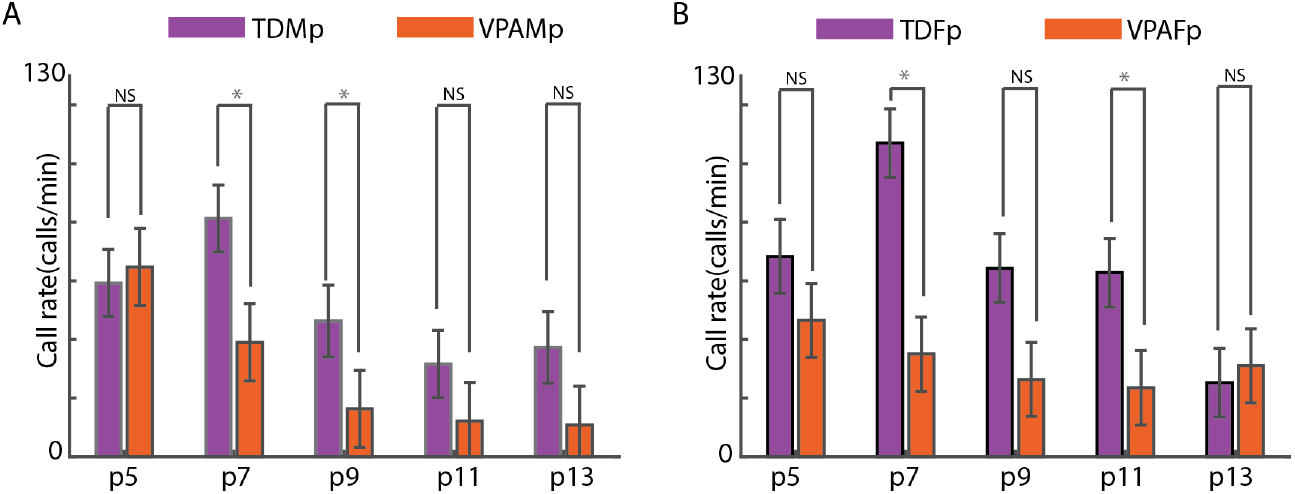
Comparative study of the call rate (calls per minute) in typically developing TD (violet) and VPA (orange) pups in A) Typically developing male pups (TDMp) versus VPA male pups (VPAMp) B) Typically developing female pups (TDFp) versus VPA female pups (VPAFp). Mann-Whitney U-test was done pairwise for each age and sex. *p<0.05, **p<0.01, ***p<0.001, NS-non significant.

### Syllable duration

The mean syllable duration for typically developing male (TDMp, violet) and VPA male (VPAMp, orange) pups is shown in Fig.3A, and similarly for typically developing female (TDFp, violet) and VPA female (VPAFp) pups in Fig.3B. On average, VPA pups have a shorter syllable duration compared to the TD (RM ANOVA). Mann-Whitney U-test pairwise for each age and sex showed a significant difference at p5 and p9 for male (p<0.05, Fig. 3A) and at p9 for females (p<0.05, Fig. 3B). A RM ANOVA reveals a significant effect of age on mean syllable duration (F (4,92) =11.02, p<0.001). Also, there is a significant interaction between group types i.e; (VPA or TD) x Sex with age (F (12,92) =2.32, p<0.05). No sex-specific difference was observed between the TD male and female group.

**Figure 3.**
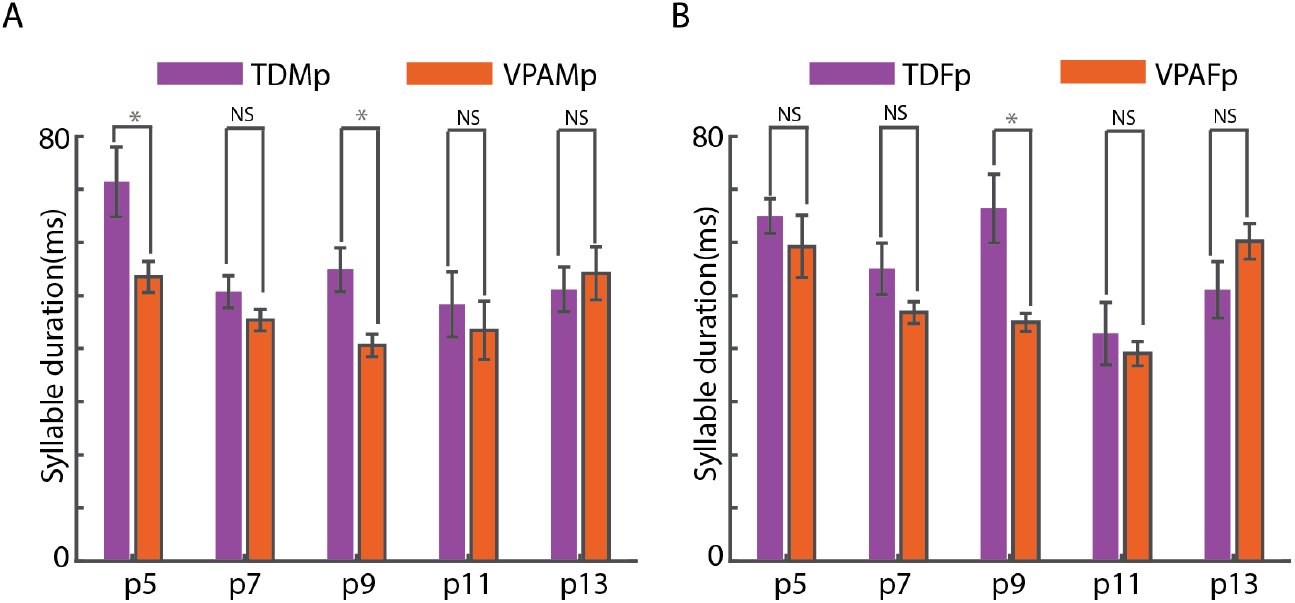
Comparative analysis of syllable duration over developmental age Comparative study of the syllable duration in A) TDMp (violet) versus VPAMp (orange) B) TDFp (violet) versus VPAFp(orange). Mann-Whitney U-test was done pairwise for each age and sex *p<0.05, **p<0.01, ***p<0.001, NS-non significant.

### Peak frequency

The peak frequency is shown for male TD (TDMp, violet) and VPA (VPAMp, orange) pups on the left side of Fig. 4 and similarly for female TD (TDFp, violet) and VPA female (VPAFp) pups on the right side). Independent of sex and age, VPA pups on average have a higher mean peak frequency than the TD (RM ANOVA). Mann-Whitney U-test pairwise for each age and sex showed a significant difference at each age for male TD versus VPA (Fig. 4A) but not for females (Fig. 4B). A RM ANOVA reveals a weak but significant effect of age on peak frequency (F (4,92) =2.6, p<0.05). There is no significant interaction between group types i.e; (VPA or TD) x Sex with age (F (12,92) =0.786, p>0.05). No sex-specific difference was observed between TDMp and TDFp.

**Figure 4.**
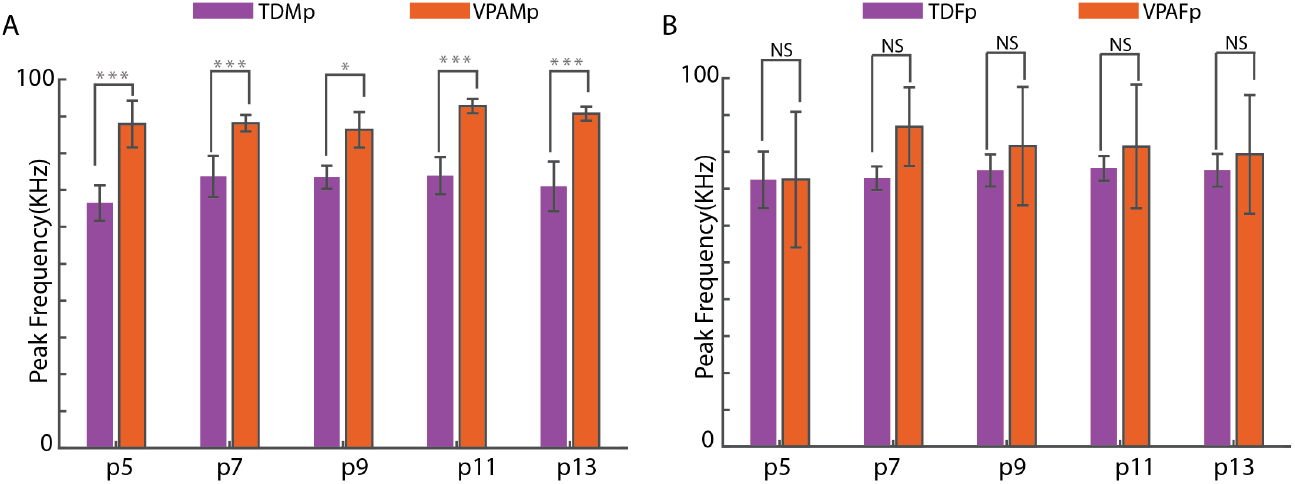
Comparative analysis of mean peak frequency over developmental age Comparative study of the mean peak frequency in A) male pups TD (TDMp,violet) versus VPAMp(orange). B) female pups TD (TDFp,violet) versus VPAFp(orange). Mann-Whitney U-test was done pairwise for each age and sex. *p<0.05, **p<0.01, ***p<0.001, NS-non significant.

Since no significant sex-specific difference was observed between TDMp and TDFp for call features, the data were pooled together and compared with pups obtained by prenatal exposure to saline. No significant difference was observed for call rate, syllable duration and mean peak frequency using RM ANOVA (Fig. S1). Therefore, the alterations observed in pup USVs could be linked to the exposure to VPA and not to any stress induced by the injection procedure during the pregnancy.

### Probability mass function (PMF) of syllables

In the current study, the syllables were categorized into four types based on their spectro-temporal features (Agarwalla et al. 2020; Agarwalla et al. 2023) as illustrated in Fig. 1B-E. To further investigate potential variations across sex and age, we examined the proportion of each syllable type in both VPA male and female pups and compared them to their respective typically developing (TD) groups. The proportion (frequency of occurrence) of syllables for TDMp and VPAMp is depicted in Fig. 5A (left top), and for TDFp and VPAFp in Fig. 5C (right top). Each row of the matrix represents the different age groups, and each column corresponds to different syllable types. For both TDMp and VPAMp, S-type syllables are dominant in proportion over age.

**Figure 5.**
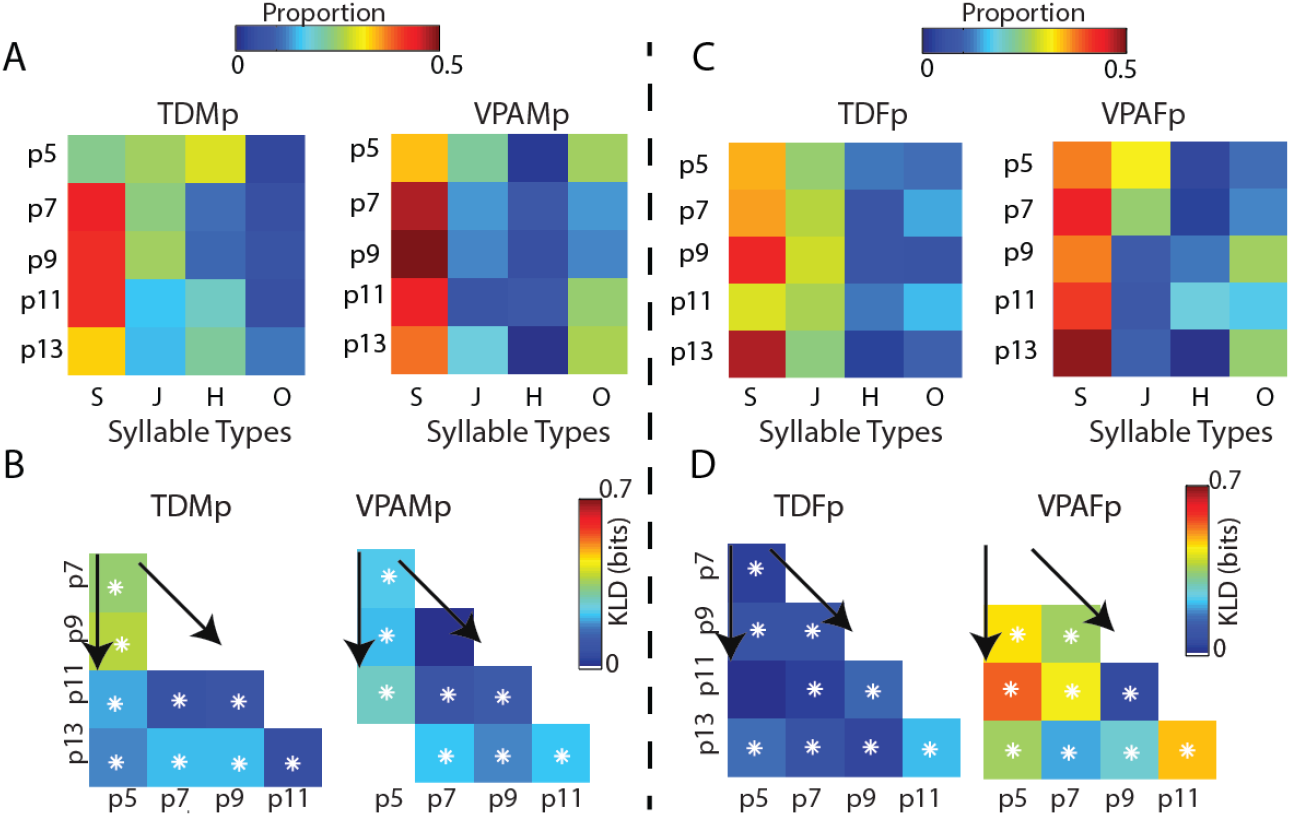
Comparison of syllable occurrence over developmental age and group The frequency of occurrence of different syllable types, termed as proportion A) typically developing male pups (TDMp, top left) and VPAMp(bottom, left). B) KLD among the distributions of typically developing male pups (TDMp, top left) and VPAMp (bottom, right) C) typically developing female pups (TDFp, top left) and VPAFp (bottom, left). D) KLD among the distributions of typically developing female pups (TDFp, top left) and VPAFp(bottom, right). White asterisk indicates significant KLD at 95% CI. The white spaces between age groups indicate a KLD of zero.

To understand how the distribution of the proportion of syllable type changes within a group with age, we first measured the Kullback Leibler Divergence (KLD) (see Methods and Materials for details) in bits for each group (TD and VPA). In Fig. 5B, the first column of the matrix, depicted by an arrow, represents how distant the distribution of the proportion of syllable types at the age of p5 are from p7(first row), p9(second row), and so on. Between any two age groups, the distance can be observed from the diagonal matrix by comparing the corresponding columns and rows of that age. In TDMp (Fig. 5B, left), the disparity in distribution diminishes with age, illustrated by the final row indicating the distance from each age to p13. The white asterisk highlights significant differences at a 95% confidence interval. Notably, there are significant discrepancies between each age in TDMp (except between p7 and p9). Conversely, in VPAMp (Fig. 5B, right), the distribution appears more consistent across ages compared to TDMp. Noteworthy the KLD distance is less between (p5, p7) and (p5,p9) in VPAMp than in TDMp.

A similar analysis was done for typically developing female pups (TDFp) and VPAFp, as shown in Fig.5D. S-type syllables are dominant in female pups as observed in male pups. In Figure 5D, the consistency in distribution across ages is more uniform in TDFp (Fig. 5D, left) than in VPAFp (Fig. 5D, right, except between p5 and p7 with no significant difference). Thus, unlike VPAMp, VPAFp exhibits more variability in the proportion of syllable types over developmental ages.

### Transition of syllables

After exploring the composition of syllables, the next step was to examine the transitions between two syllables (see Methods and Materials for details). Sequences of syllables were analyzed considering two successive syllables, considering only the starting two syllables in bouts. Syllable-to-syllable transitions were quantified using the joint probability distributions, which represent the probability of occurrence of every possible pair of syllable types. The joint distributions are displayed in the form of matrices in Fig.6A (for the first two syllables of a bout) for TDMp (first column) and VPAMp (second column) across developmental age. For TDMp, S-S, S-J, J-J, J-S, and H-H transitions are the most prominent across all the ages except p13, where J-J and J-S transitions are insignificant. In contrast, VPAMp has no H-H transitions and exhibits changes with age without a consistent pattern. Additionally, the J-S, J-J were not significant in VPAMp after p5. The diagonal matrices in Fig.6B show the KLD among the distributions.

A similar analysis was done for syllable-to-syllable transition for females Fig. 6C. TDFp (third column) showed a consistent pattern of transitions, including S-S, S-J, J-S, and J-J across all the ages. However, for VPAFp (Fig. 6C, fourth column), there was considerable variability in the pattern with age. Similar to VPAMp even in VPAFp, the H-H, J-J, and J-S (except p5) transitions were missing. Interestingly, the H-H transitions were present only in TDMp. The KLD among the distributions is shown by the diagonal matrices in Fig.6D Thus, TD groups, independent of sex, maintained a pattern that was altered in VPA pups.

**Figure 6.**
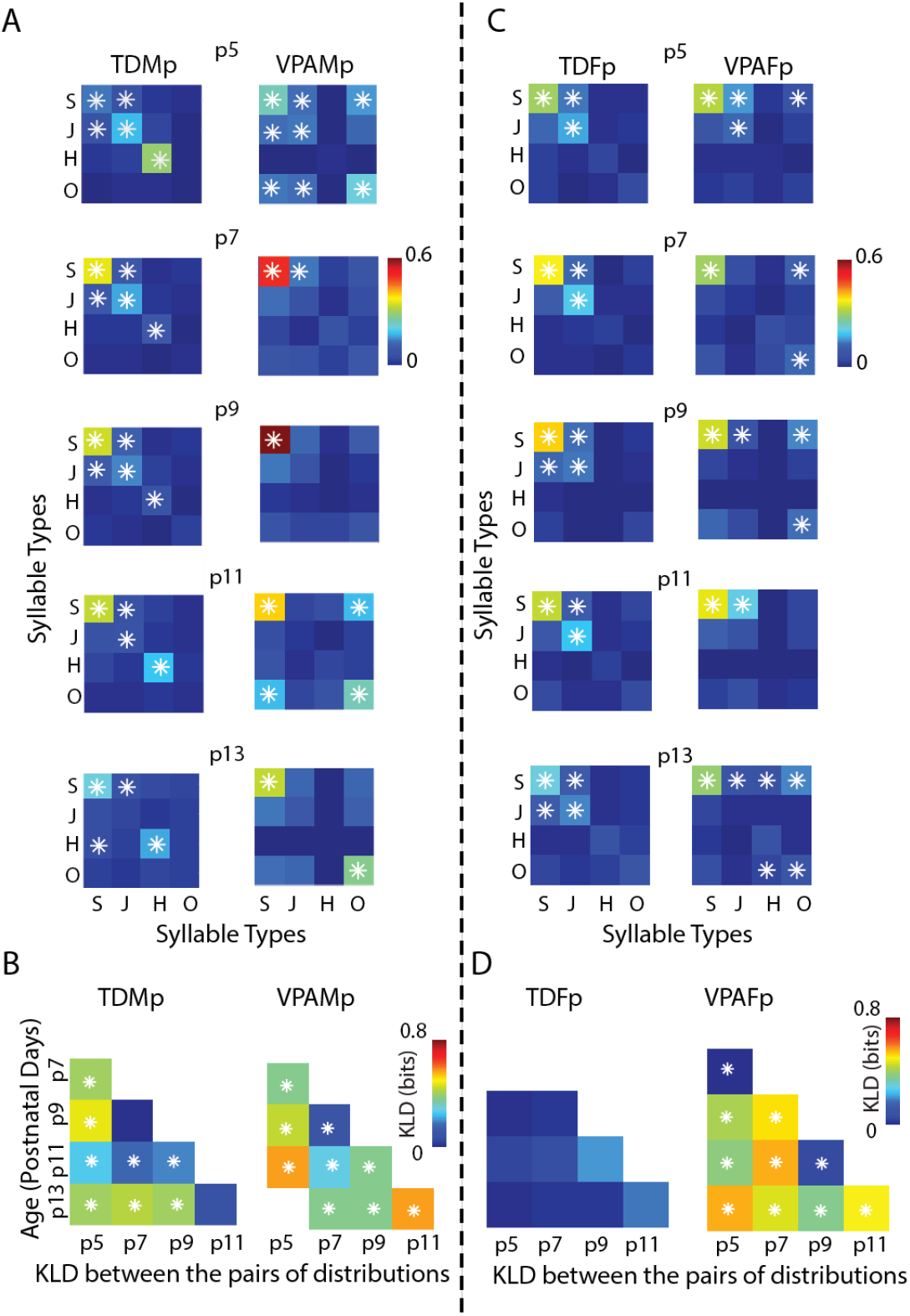
Syllable to syllable transition probability matrices. (A) Joint probability distributions of syllable type to syllable type transition considering starting two syllables in bouts is depicted in each of five matrices for each age p5, p7, p9, p11 and p13 for the two groups of pups, TDMp (first column) and VPAMp (second column) (Asterisks indicate significance at 95% confidence). (B) The diagonal matrix plot quantifies the *KLD* between joint distributions for TDMp (left) and VPAMp (right). (C)Joint probability distributions of syllable type to syllable type transition considering starting two syllables in bouts is depicted in each of five matrices for each age p5, p7, p9, p11 and p13 for the two groups of pups, TDFp (first column) and VPAFp (second column) (Asterisks indicate significance at 95% confidence). (D) The diagonal matrix plot quantifies the *KLD* between joint distributions for TDFp (left) and VPAFp(right).

### Differences between male and female in typically developing vs. VPA pups: Transition vs. Single Syllable Levels

Females have a higher ability to camouflage their autistic symptoms compared to males (Dean et al., 2017; Dworzynski et al., 2012). Therefore, we investigated whether the differences between TD and VPA pups are more pronounced in males than in females across developmental ages. We calculated the Kullback-Leibler Divergence (KLD) between TD and VPA groups, separately for male and female pups.

In Fig. 7A, the distance between the proportions of syllable types between TD and VPA pups for each age group is shown for males (solid lines) and females (dashed lines). Significant differences were observed at all ages at the 95% CI between the TDMp and VPAMp distributions of syllable proportions, with the highest difference noted at p5 (solid lines). Similarly, the distribution of syllable type proportions between TDFp and VPAFp was significantly different at each age, except at p5 and p7, with the highest difference at p9 (dashed line). Interestingly, differences between TD and VPA pups were greater in males than in females at p5 and p11, while females exhibited the highest differences at p9. At p7 and p13, differences between sexes were comparable.

**Figure 7.**
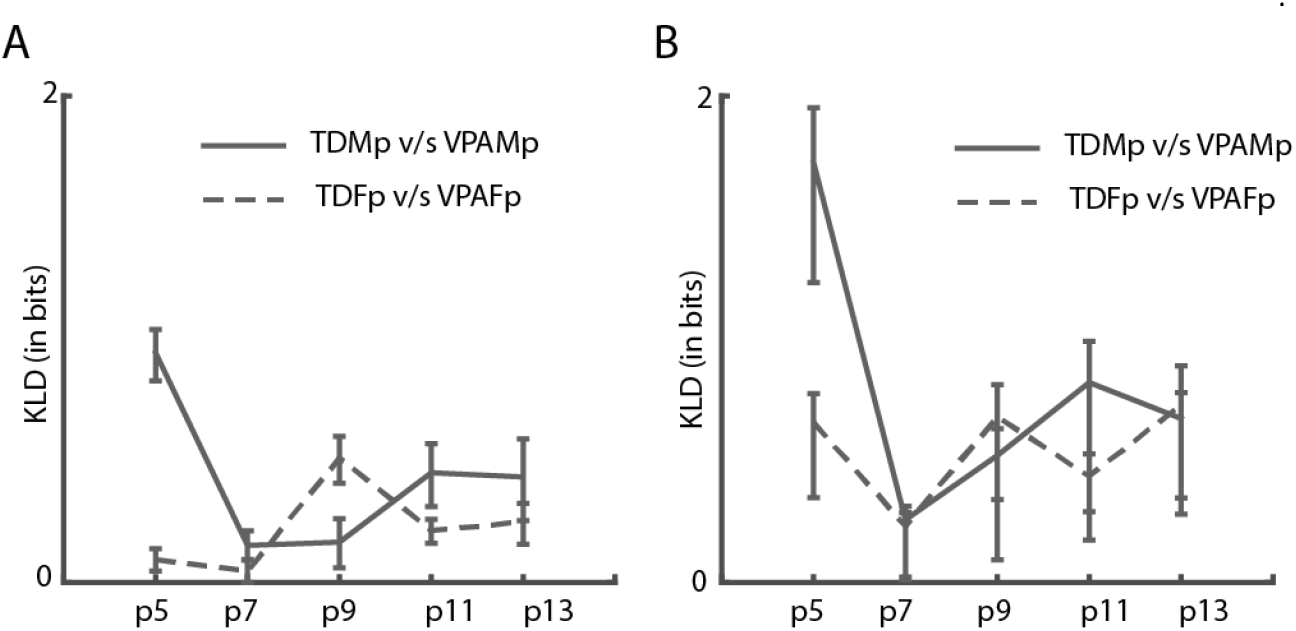
Comparative KLD Analysis of Syllable Proportions and Transitions in Male and Female Pups A) KLD (in bits) between the distribution of the proportion of syllable types between TDMp and VPAMp shown in solid black lines; TDFp and VPAFp in dashed black lines for each age. B) KLD (in bits) between TDMp and VPAMp joint distributions; TDFp and VPAFp shown in solid black lines; TDFp and VPAFp in dashed black lines for each age. The error bars indicate 95 % CI.

Additionally, the KLD between syllable transitions was quantified (Fig. 7B). A significant difference was observed between TDMp and VPAMp at each age, with the highest difference at p5, similar to the syllable proportions, but more pronounced. In contrast to the proportion of syllables, differences in syllable transitions between TD and VPA pups were comparable between males and females at all other ages.

### Information theoretic measures reveal lack of structure in VPA pups independent of sex

To check for the presence of higher-order structure in the sequences of syllables first, we first investigated if there was any dependence between syllables. Essentially, we probed to see if there was any order in the occurrence of the syllables or if they occur randomly. To test the same, we computed *Mutual Information (MI)* between the syllable at the start of a bout and the syllable at subsequent positions of a bout (Agarwalla et al. 2020; Agarwalla et al. 2023). Significant dependence or *MI* between the syllables treated as random variables would indicate the presence of structure in the sequence, while lack of dependence or ‘0’ *MI* would indicate a lack of structure. Fig. 8 and 9 show the debiased *MI* between the first syllable and the *n*th (*n*=2, 3, …) with 95% confidence intervals for all the groups (solid line). The baseline for the *MI* is represented with 95% confidence intervals for the scrambled sequences, indicating the *MI* for no dependence (dashed gray line). *MI* estimates with no overlap between the two confidence intervals at each position were considered significant. There is a clear presence of structure in TDMp (Fig.8A, top) and TDFp (Fig.9A, top) across all ages. However, given the first syllable, the positions to which it can be predicted vary. In VPAMp (Fig. 8B, bottom) and VPAFp (Fig. 9B, bottom), we observe either no or less dependency compared to TD groups. This suggests that VPA pups lack predictive-ness among the syllables, which is well captured in typically developing pups across different developmental ages. Additionally, we tracked the informative sequences (see Methods and Materials section) as tabulated in Table 2. The sequences with S-S transitions were common to all the groups. The TD group of males and females also exhibited sequences with J-J and J-S transitions, which were missing in VPA pups.

**Table 2:** Informative sequences in each group over developmental age. The sequences tracked are represented by X(N), where X denotes the syllable type and N indicates the number of times the X syllable is repeated in the series (X-X-X… N times). The S-type (S), Jump-type (J), Harmonic (H), Other (O) are indicated by the acronyms as shown in brackets corresponding to each syllable type.

|  | TDMp | VPAMp | TDFp | VPAFp |
| --- | --- | --- | --- | --- |
| **p5** | S(4)<br>J(4) | S(4) | S(5)<br>J(4) | S(4) |
| **p7** | S(7)<br>J S(2)<br>J(2) S | S(5) | S(6)<br>J(2) S<br>J(3) | S(3) |
| **p9** | S(5)<br>J(3) | S(3) | S(5)<br>J(4) | S(4) |
| **p11** | S(4) | S(2) | S(4)<br>J(3) | S(4) |
| **p13** | S(4) | S(2) | S(3)<br>J(2) | S(4) |

**Figure 8.**
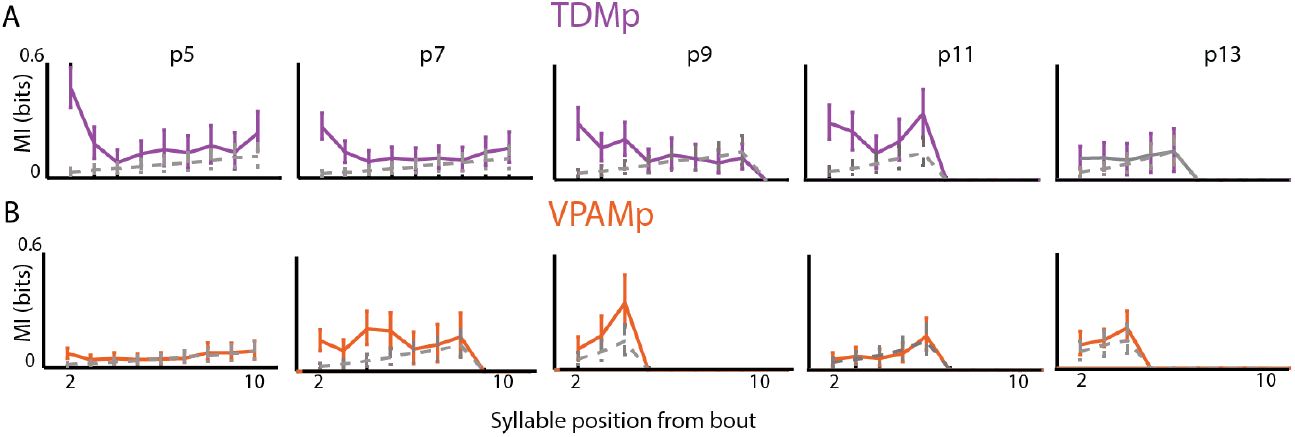
*MI* between the first and the *n*th syllable from the starting of a bout for male pups. The four plots show the dependence (quantified by *MI* in bits) between the first syllable and the subsequent syllables in the A) TDMp (violet) and B) VPAMp (orange). The actual *MI* is plotted in solid lines with error bars showing 95% confidence intervals. The dashed thin gray line shows the *MI* (with 95% confidence intervals) between the syllables at the same positions when the syllable sequences in bouts were randomly scrambled.

**Figure 9.**
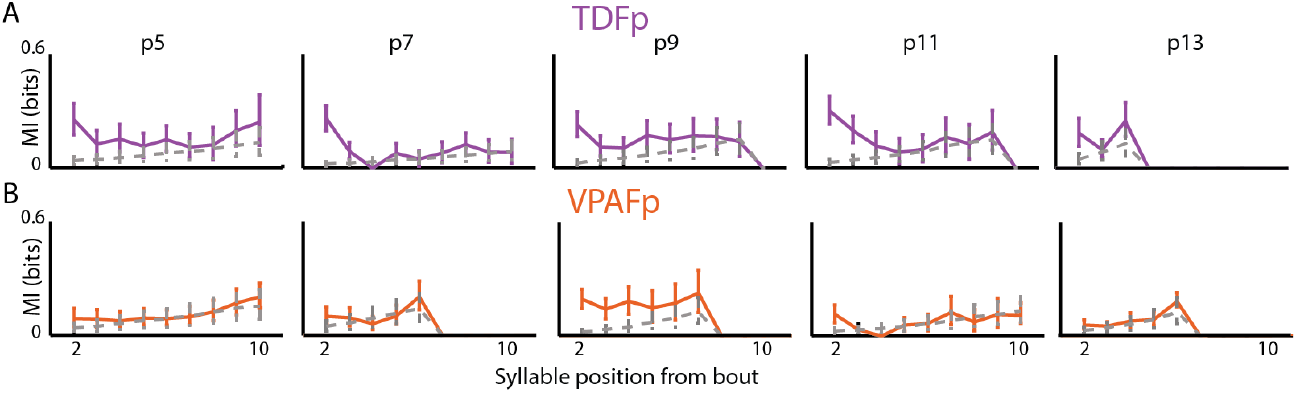
MI between the first and the *n*th syllable from the starting of a bout for female pups. The figure is arranged in the same way as Fig. 8, with data from female pups A) TDFp (violet) B) VPAFp (orange).

## Discussion

In the current study, we leveraged the predictability and structure in the mouse USVs (Agarwalla et al. 2020; Agarwalla et al. 2023; Grimsley et al. 2011), particularly in PICs, along with acoustic features, as an assay to probe ASD-like phenotypes in a VPA mouse model. We observed a general reduction in call rate, elevated peak frequency, and shorter call duration in VPA pups. However, these acoustic features displayed considerable diversity across developmental ages (p5-p13) and sex, with significant differences observed at certain ages/sexes while not evident at others. As an illustration, the peak frequency exhibited changes in VPAMp compared to the TD group across all ages. However, in VPAFp, this alteration was not noticeable at any age, suggesting that female pups either showed signs of recovery or were less impacted compared to males. Whereas for call rate and syllable duration at some ages for both sex significant differences were observed. However, the alteration in the VPA pups for the transition of syllables, the predictability, quantified using *MI*, and the informative sequences emitted over developmental age, were observed in both sexes and across all ages. Therefore, utilizing higher-order vocal structures along with call features in mouse USVs appears to be a more promising measure for interpreting abnormalities related to social communication deficits.

VPA mouse is a validated model and can serve as a valuable tool to investigate the neurobiology underlying ASD-like behavior and test novel therapeutics (Moore et al. 2000; Williams et al. 2001; Rasalam et al. 2005; Rodier et al. 1996). The neonatal mouse pup USVs being sensitive to social context can serve as a proxy for reduced social attachment or atypical social information processing. Consequently, various studies to date have investigated the effects on USVs emitted by the pups due to the unavailability of easily measurable behavioral indices (Semple et al. 2013; Silverman et al. 2010), aiming for early detection (Vivanti et al. 2014) to enable timely interventions (Okoye et al. 2023). The USV acoustic features and the call sequences emitted undergo modulation with age in typically developing pups (Grimsley et al. 2011), tracking the same in the VPA pups to monitor any alterations is essential. Despite targeting specific postnatal ages and sexes, as well as pooled data across different ages/sexes, and monitoring developmental changes in both sexes, the changes in mouse USV call features over sex and age remains elusive. For instance, Gandal et al. 2010 observed decreased call rates in combined mouse samples across p2, p5, p8, and p12. Moldrich et al. 2013 and Cheaha et al. 2015, examined only male mice at p8 and p12, respectively, noting a decreased call rate. Kuo et al. 2017, also focusing on male mice at p8, reported reduced call rates, and other features such as decreased duration and peak frequency. Tartaglione et al. 2019 considered mice of both sexes from p4 to p12; at p10, females exhibited shorter call durations, but no other postnatal days showed significant effects. Tsuji et al. 2020, studying mice of both sexes from p3 to p14, found no differences in peak frequency in either sex and noted lower call rates in both sexes at p11. Thus, a comprehensive understanding of how USV features change across development and between sexes still remains unclear. In our current study, we found on average a decrease in call rate (Fig. 2), increased peak frequency (Fig. 3), and shorter call duration (Fig. 4) for VPA pups compared to TD pups which varied across ages and sex. The call rate was significantly reduced in VPAMp at p7 and p9 whereas in VPAFp, the effect was significant at p7 and p11 compared to typically developing ones. Syllable duration was shorter for VPAMp at p5 and p9 and for VPAFp at p9. The peak frequency was higher for VPAMp at all ages but not female. There seems to be a lot of variability in the basic call features over sex and age.

Alteration in the syllable composition was observed by Cheaha et al. 2015(mice), Moldrich et al. 2013 at p8(mice); Tsuiji et al., 2021(mice), p11. However, none of the studies to the best of our knowledge have looked at the composition of syllables by monitoring over developmental age and also exploring the within group (say TDMp over age) variability as a function of age for both sexes. The frequency of occurrence of syllable types (Figure 5 A, C) was altered in both VPAMp and VPAFp. Interestingly, the within-group variability in the composition of syllables was higher in VPAFp over age than in any other group (Figure 5B, D), an exciting pattern unique to VPA females. Thus, the way ASDs manifests in female mice might be different from males for a few features. At the syllable transitions level too alterations were observed in both male and female VPA pups compared to TD groups (Figure 6A, C), with more pronounced age-related changes within the VPA female group compared to typically developing female pups (TDFp), as quantified using KLD (Figure 6B, D). Additionally, certain transitions, such as J-J (except at p5) and H-H, were consistently present only within the TDM group, a pattern that held true even when analyzed within individual litters, highlighting sex-specific differences at the transition level.

Previous clinical studies in humans have shown that females often have a greater capacity to mask their autistic symptoms compared to males (Dean et al., 2017; Dworzynski et al., 2012). To explore this, we examined whether differences between TD and VPA pups are more prominent in males than in females across various developmental stages. By quantifying the disparity between the VPA and typically developing pups at each age using KLD, we found that the differences were heightened in TD and VPA males compared to females at p5, both at the syllable composition (Figure 7A) and transition level (Figure 7B). This indicates that at p5, which marks the onset of vocalization, the differences in TD and VPA males are more prominent than females at any other age.

Certain studies (Agarwalla et al. 2020; Agarwalla et al. 2023; Grimsley et al. 2011) have highlighted the presence of predictability in mouse USV sequences, akin to human speech and songbirds at the rudimentary level and tracked the sequences with high predictability, named as informative sequences. The tracked sequences have been proposed to carry behavioral significance beyond mere mathematical outcomes (Agarwalla et al. 2023). Agarwalla et al. 2020, reported lack of MI and altered sequences in a genetic ASD mouse model at p5, alterations being specific only for male pups and not in females. The same has not been looked into for VPA pups. We thereby explored the possibility of alterations in structure and the speech-like sequential predictability (Agarwalla et al. 2020; Agarwalla et al. 2023) of PICs. To further investigate the predictability present in the USV sequences, MI was used to measure the dependency of the preceding syllables on the syllable considered to be the starting of a sequence. Typically developing pups had significant MI across all ages (except p13), whereas the VPA group lacked it (except minor dependencies at p7, VPAMp, and p9, VPAFp). We also tracked the informative sequences emitted by each group and found that sequences with S(N) transitions were common to all groups, although their lengths (N) varied across sex and age. However, the informative sequences with J(N) or JS(N) transitions (length, N varied) which were present in TD groups, were absent in VPA groups (Table 2). Thus, VPA pups lacked complex higher-order call structures present in TD. Understanding sex- and age-specific features, as well as independent characteristics in ASD research, is crucial for uncovering the core aspects of the disorder, identifying biomarkers, and developing effective treatments. By focusing on features that transcend sex and age differences, researchers can enhance our understanding of ASD and improve outcomes for individuals affected by the condition.

While the current study provides insights into changes in ultrasonic vocalization features and higher-order vocalization structures in VPA pups compared to typically developing ones across developmental ages, both for males and females, the circuit-level alterations that could mechanistically explain these findings remain unexplored. For instance, a specialized group of neurons in the midbrain periaqueductal gray (PAG-USV neurons) have been shown to act as a mandatory gate for ultrasonic vocalization (USV) production in adult mice (Michael et al., 2020). But how the circuitry changes over developmental age in typically developing and VPA pups would help envision better which pathway or circuitry is altered and if could be utilized to explain any sex-specific changes. Additionally, the differences between VPA and TD pups might be due to alterations in the early neural circuitry in Layer V neurons in the motor cortex, which are active during singing, project directly to brainstem vocal motor neurons, and are necessary for keeping song more stereotyped and on pitch (Arriaga et al., 2012). While the excitation-inhibition imbalance (Rubenstein and Merzenich, 2003; Gao and Penzes 2015) in the neural circuits is a well-established phenomenon observed in ASD models, how these factors impact vocalization and the interplay between excitatory and different inhibitory interneurons subtypes like parvalbumin, somatostatin etc requires further investigation. Further research in these areas could provide insights and hypotheses about the circuit-level alterations that might mechanistically explain the findings in our study.

## Supplementary Figure

**Figure S1:**
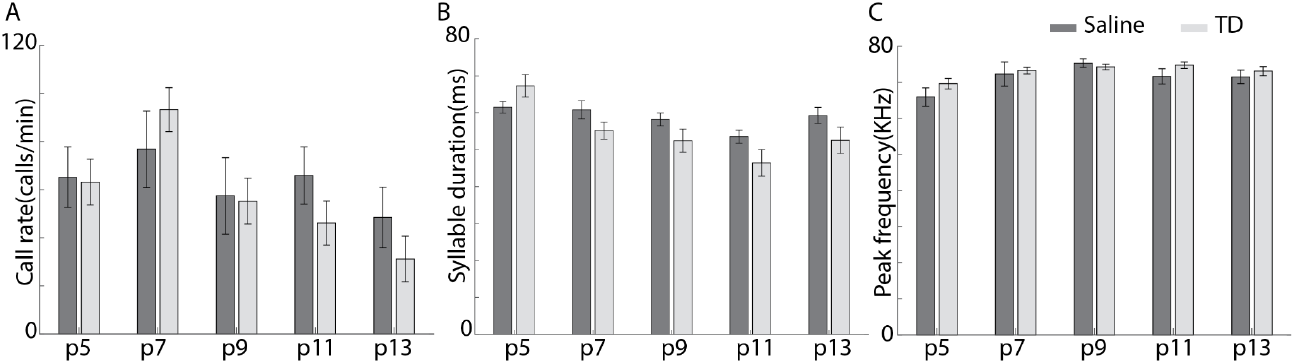
Comparative analysis of acoustic features over developmental age between Saline and TD. Comparative study of the A) Call rate B) Syllable duration distribution C) Mean peak frequency Repeated measure ANOVA with Post-hoc analysis by Tukey multiple comparisons. No significant difference observed for the acoustic features.

